# Age and maturation stage linked consequences of fibrinogen on human oligodendroglia

**DOI:** 10.1101/2024.05.27.596068

**Authors:** Gabriela J. Blaszczyk, Chao Weng, Abdulshakour Mohammadnia, Qiao-Ling Cui, Arianna Giurleo, Adam M.R. Groh, Chloe Plouffe, Julien Sirois, Valerio E. C. Piscopo, Moein Yaqubi, Asad Taqvi, Erin Cassidy, Liam Callahan Martin, Jeffery A. Hall, Roy W.R. Dudley, Myriam Srour, Stephanie E J Zandee, Wendy Klement, Sandra Larouche, Alexandre Prat, Thomas M. Durcan, Jo Anne Stratton, Jack P. Antel, G. R. Wayne Moore

## Abstract

Fibrinogen is a blood-derived protein involved in coagulation, and can make its way into the central nervous system (CNS) following breakdown of the blood-brain barrier. This molecule has been implicated in multiple sclerosis (MS), a disease marked by inflammation and demyelination in the CNS, as well as other neurological disorders. However, the effect of this molecule has not been studied on human myelinating cells. This study examines how fibrinogen influences human oligodendrocyte (OL) lineage cells at various stages of development. Using induced pluripotent stem cell-derived (iPSC) OL precursors and human primary OLs, we examined the effects of fibrinogen on cell differentiation, viability and myelination-related function. Here we show that fibrinogen induces an aberrant differentiation of early lineage OLs, by inhibiting their maturation and inducing an astrocytic phenotype, as seen in previous studies. On mature OLs, fibrinogen was found to promote myelination capacity as shown by ensheathment assays as well as on the RNA level. These effects were associated with the activation of BMP signalling, both in early and mature OLs. Transcriptomic analysis of human MS brain tissue shows similar pro-myelination changes in a subset of OLs, suggesting in vivo relevance. These findings indicate that fibrinogen has a lineage-dependent effect, where it may be inhibitory earlier in the lineage while promoting OL function in later stages. Understanding this dual role will provide insight into remyelination failure in MS and highlights the importance of timing and target in future therapeutic strategies.

**Significance Statement:** In multiple sclerosis (MS), the blood protein fibrinogen leaks into the brain and has been shown to interfere with myelin repair. This study demonstrates that fibrinogen has opposite effects on human oligodendrocyte-lineage cells depending on their stage of maturation. While it blocks the differentiation of early-stage cells, it enhances the functional capacity of mature oligodendrocytes. These findings help explain why remyelination may fail in MS and suggest that fibrinogen could both hinder and support repair, depending on the cell context. This dual role has important implications for developing stage-specific therapies for MS.

## 1. Introduction

Potential contributors to the injury and repair responses of oligodendroglia (OLs) are blood-derived factors that may access the central nervous system (CNS) due to the breakdown of the blood-brain barrier (BBB) (1). Fibrinogen is a circulating blood clotting factor composed of polypeptide chains which have polymerization sites for clot formation (2), and various domains known to be involved in stimulation of multiple cell types (3). Fibrinogen has been described in multiple sclerosis (MS) and detected in the borders of chronic active white matter plaques (4) and leakage into diffusely abnormal white matter (5) and normal-appearing white matter (6). It has also been found in MS motor cortex, where its presence significantly correlated with reduced neuronal density (7), implicating its direct role in cellular damage. Moreover, plasma levels of fibrinogen are elevated in MS relapses (8), and its role in MS pathogenesis has been linked to its function in the coagulation cascade and as a pro-inflammatory agent (9). The role of fibrinogen in MS and autoimmune disorders of the CNS has been extensively studied (10), and rodent studies have shown that fibrinogen can inhibit the differentiation of rodent OPCs into myelinating oligodendrocytes, primarily mediated via bone morphogenetic protein (BMP) signaling (11). While fibrinogen is thought to have an important role in neuroinflammation (10, 12) and the pathogenesis of MS (9, 13-16), it is unknown if these findings are translatable to human oligodendroglia across their development.

The objectives of our study were to extend insights into the direct effects of fibrinogen on human OLs in relation to age and lineage stage. We employed primary human mature OLs (O4+A2B5-) and late lineage cells (O4+A2B5+) (17, 18) derived from surgically-resected tissue samples together with early-stage progenitor cells derived from human induced pluripotential stem cells (iPSCs), as such primary cells cannot currently be derived from available human CNS sources. Primary cell studies were also considered in context of the donor age, as we previously reported that ensheathment capacity, as determined using cultures with synthetic nanofibers, and susceptibility to injury were donor age dependent (19). For iPSC studies, we also employed a previously characterized reporter line, where the degree of expression correlated with the differentiation stage of the cells (20). Using these methods, we found that adult-derived mature OLs respond to fibrinogen by increasing their ensheathment capacity. While pediatric late-stage progenitors, and iPSC-derived early progenitors decrease their differentiation, in line with previous reports that studied various other toxic effects on OLs.

## 2. Materials and Methods

This study was ethically approved by the Institutional Review Board of the McGill University Faculty of Medicine and Health Sciences (A09-M72-22B [22-08-053]).

### Antibodies

Antibodies used in this study are listed in Table S3.

### Human primary cell isolation and culture

Studies were conducted on OLs derived from pediatric (2.5 -18 years) and adult (26 - 65 years) tissue samples collected from patients undergoing surgery, regardless of biological sex. The use of adult tissues was approved by the Montreal Neurological Institute Neurosciences Research Ethics Board and the use of paediatric tissues by the Montreal Children’s Hospital Research Ethics Board. As previously described (18, 19), samples were dissociated by trypsin digestion and subjected to Percoll gradient (Sigma, Oakville, ON) separation. To enrich progenitor populations, MACS magnetic beads selection against the cell surface marker A2B5 (Miltenyi Biotec, Auburn, CA) was performed following cell isolation, and positive and negative fractions were cultured as described previously (18, 19).

Our previous detailed flow cytometry based analysis of the adult and fetal cell populations (21) indicated that of the oligodendroglial (OL) population isolated from the adult brain 5-10% of the cells were recognized by the A2B5 antibody (referred to as A2B5+ cells), as also previously reported by others (22), and are considered late progenitor cells. 85-90% of these cells express detectible levels of O4. As previously documented, these cell preparations are devoid of astrocytes and neurons and have a small proportion of microglia as documented by FACS (21) and single cell mRNA sequencing (Supp Fig 1 in Luo et al (19)). The two cell fractions (A2B5+ and A2B5-) express comparable levels of mature myelin genes (MBP and PLP) based on bulk RNA sequencing whereas progenitor genes (PDGFRa and PTPRZ1) are more highly expressed in the A2B5+ cells (Fig 3 in Luo et al (19)).

**Figure 1.**
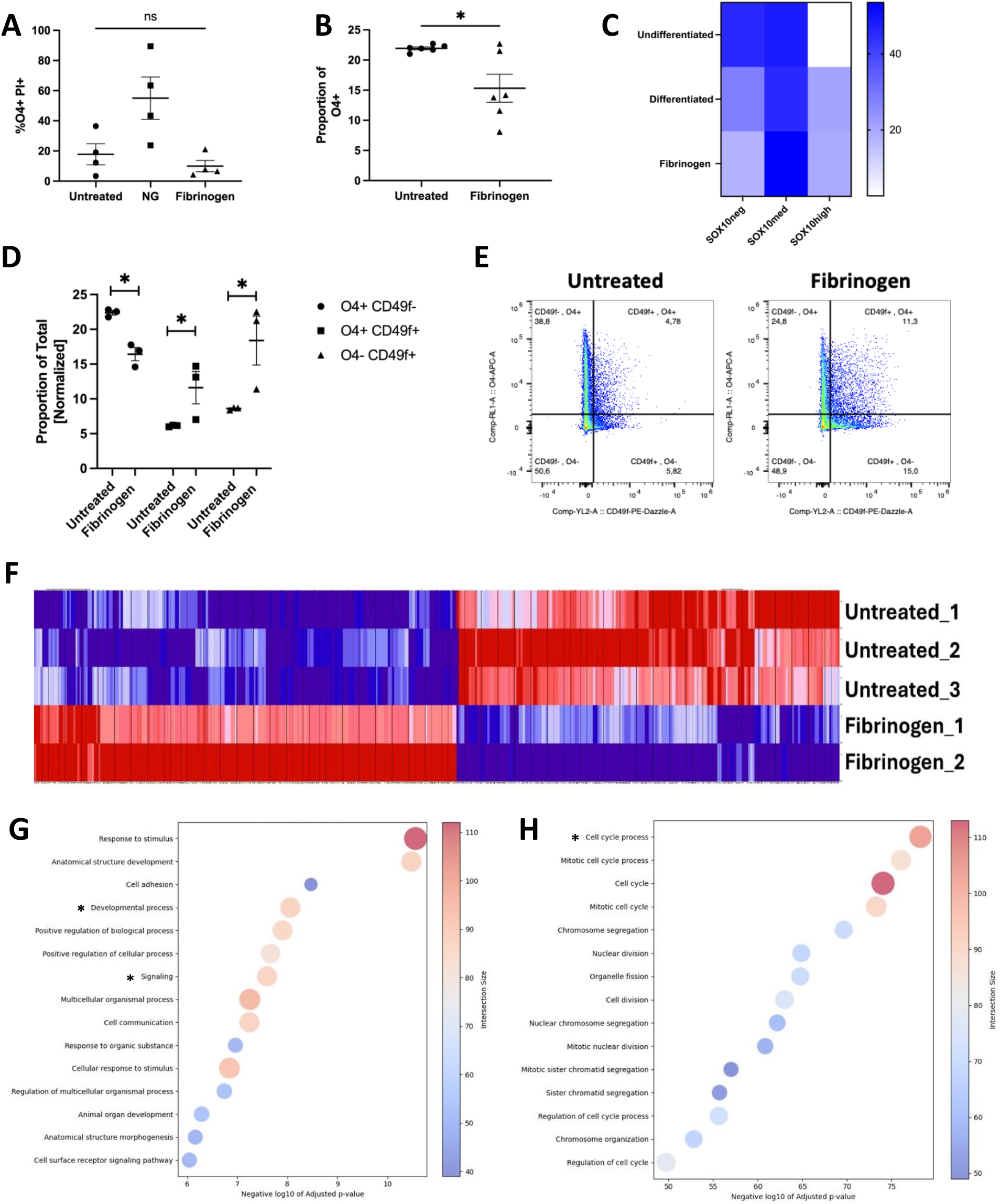
Fibrinogen effect on human iPSC-derived OPCs. **A**. Proportion of viable OPCs (O4+) following immunofluorescence staining for propidium iodide (PI). NG: no glucose. Bars indicate SEM. **B**. Proportion of OPCs (O4+) as quantified following immunofluorescence microscopy, showing a decrease in OPCs following fibrinogen treatment for 4 days. Bars indicate SEM. * p < 0.05 **C**. Proportions (%) of iPSC-derived cultures expressing SOX10mO as measured by flow cytometry, average of n=5 passages. **D**. Proportions of live cells co-expressing OPC/astrocyte markers as measured by flow cytometry. Values normalized to the average of the untreated control, n=3 passages. Bars indicate SEM. * p < 0.05. Paired t-tests were applied to assess significance levels in all plots. **E**. Flow cytometry gating technique. **F**. Differentially expressed genes following bulk RNAseq analysis of sorted SOX10mO-med cells. **G-H**. GO term analysis of (G) downregulated and (H) upregulated genes.

#### Viability assays in dissociated cell cultures

The cells were cultured in DMEM/F12+ N1 medium + 0.1% BSA + Pen/Strep + B27 medium (Life Technologies, Grand Island, NY) in 96 well plates (Fisher Scientific) and treated with plasminogen-depleted fibrinogen (Millipore Sigma, Burlington MA, 341578, 80ug/ml to 4mg/ml) for 6 days, with addition of fibrinogen alongside media changes every other day. Cells were stained live for O4 (1:200, R&D systems, Oakville, ON) and the cell viability was determined by live staining with propidium iodide (PI, 1:200, Thermo Fisher Scientific, Mississauga, ON).

### Nanofiber ensheathment cultures

Aliquots of A2B5+ and A2B5-cells were plated in multi-well aligned poly-L lactide nanofiber plates (The Electrospinning Company Ltd., Didcot, Oxfordshire, UK) (10,000 cells per well) and treated with plasminogen-depleted fibrinogen (2.5 mg/ml) in DMEM/F12+ N1 medium supplemented with B27 (Life Technologies, Grand Island, NY) and T3 (Sigma-Aldrich, Oakville, ON) for two weeks. Fibrinogen was supplemented alongside media changes every other day. Cell cultures were immunolabelled with O4 antibody diluted 1:200 (R&D Systems, Oakville, ON) for one hour, followed by corresponding secondary antibody (1:500) conjugated with Alexa Fluor 647 (Thermo Fisher Scientific, Mississauga, ON) for two hours. To determine the percentage of O4+ cells ensheathing nanofibers in a culture, a blinded rater counted individual O4+ cells whose nanofibre-ensheathing processes could be identified as being connected to their cell bodies identified by Hoechst 33342 (Thermo Fisher Scientific, Mississauga, ON) staining of nuclei. Ensheathed processes were recognized by their straight alignment with underlying nanofibers and increased thickness of the segments (Figure 1F). Cells were assigned as ensheathing (one or more segments) or not and counted by the Plugin Cell Counter ImageJ program.

### Human iPSC culture and OPC differentiation

iPSC-derived OL lineage cells were differentiated according to a recently published protocol (20). The use of iPSCs and stem cells in this research was approved by the McGill University Health Centre Research Ethics Board (DURCAN_IPSC/2019–5374). iPSCs were used from healthy control lines as well as the SOX10-mOrange reporter line (20), which were generated at the Montreal Neurological Institute’s (MNI) Early Drug Discovery Unit (EDDU) (Table S2). According to a previously published protocol (20), early OL-lineage progenitors were generated, and plated on poly-L-ornithine (Sigma-Aldrich, Oakville ON)/ laminin (Invitrogen, Waltham MA) coated culture vessels. Post day 75 *in vitro*, cells are used for up to a maximum of four passages and differentiated to reach a majority of OL-lineage cells at a total of 21 days. As regards the composition of iPSC derived cells, at post day 75 *in vitro* (oligodendrocyte precursor cell [OPC] phase) ∼50% of the cells express the early marker PDGFRα and ∼36% the ganglioside A2B5. Between 20% and 40% of the total cells express O4 (20).

Cells were treated with plasminogen-depleted fibrinogen (Millipore Sigma, Burlington MA, 341578, 2.5 mg/ml) for 4 days prior to fixation (days 18-21 of differentiation, in differentiation medium). The cell viability (PI) and differentiation (O4) were determined as described above. Proliferation was determined by staining with Ki67 (Invitrogen, Waltham MA), post-fixation and permeabilization diluted 1:200, overnight. Secondary antibody (Alexa Fluor 488 or 647; Invitrogen, Waltham MA) was applied the following day, diluted to 1:400.

### Immunofluorescence Analysis

Plates were imaged with a 10X objective using a Zeiss Axio Observer fluorescence microscope (Carl Zeiss Canada, Toronto) or the ImageXpress (Molecular Devices, San Jose, CA) high-content imaging platform following staining for O4, PI or Ki67. Cells were counted by a blinded individual using the ImageJ software or automatically using the ImageXpress software.

### Signaling assessment

#### Staining

Primary human OLs and iPSC-derived OPCs (following 21 days of differentiation) were treated with molecules for 2 hours to induce signaling. The following concentrations were applied: plasminogen-depleted fibrinogen (Millipore Sigma, Burlington MA, 341578, 2.5 mg/ml), recombinant human BMP4 (Peprotech, Rocky Hill NJ, 50 ng/mL), BMP4-inhibitor DMH1 (Selleck Chemicals, Houston TX, 2 uM). Post-fixation, cells were permeabilized with 100% cold methanol and stained with anti-phosphorylated (p)SMAD1/5/9 (Cell Signaling Technology, Danvers MA) overnight at a 1:50 dilution, and secondary antibodies (Alexa Fluor 488, 555 or 647; Invitrogen, Waltham MA) diluted to 1:400 were added for two hours the following day.

#### To measure Mean Fluorescence Intensity (MFI)

Images were acquired on a Zeiss Axio Observer fluorescence microscope (Carl Zeiss Canada, Toronto, ON) at 20X objective or the Zeiss LSM 900 confocal microscope at 20X objective. For pSMAD1/5/9 nuclear intensity, nuclei of O4+ cells were selected by a blinded individual on the ImageJ software and the mean grey value of the pSMAD1/5/9 channel was recorded.

#### To record pSMAD localization

Following image acquisition at the 20X objective, localization of pSMAD1/5/9 on O4+ cells was assessed by a blinded individual based on four criteria: no signal, cytoplasm only, nucleus and cytoplasm, nuclear only. 20-25 cells were counted for each replicate.

### FACS Sorting and Flow Cytometry

Using the iPSC-reporter line SOX10mO (20), cells were collected from the culture vessel using TrypLE (Thermo-Fisher, Mississauga, ON) added for 30 mins at 37□. The cells were passed through a 40µM mesh to ensure a single cell suspension. Following resuspension in PBS (Wisent, Saint-Jean-Baptiste, QC), the cells were counted using the Attune flow cytometer (Thermo-Fisher Scientific, Mississauga, ON). Sorting was based on reporter intensity using the previously described protocol (20), with the Aria fusion cell sorter (BD Biosciences, Franklin Lakes NJ). For phenotyping, cells were stained as previously described (20), and acquired on the Attune flow cytometer (Thermo-Fisher Scientific, Mississauga, ON). Data was analyzed using the FlowJo software (Ashland, OR).

### Bulk RNAseq preparation and analysis

Three subsequent passages of FACS-sorted SOX10-mOrange cells were collected as replicates, and primary cells from two different donors (53 yr, 58 yr) were collected as replicates following treatment with plasminogen-depleted fibrinogen (Millipore Sigma, Burlington MA, 2.5 mg/ml) for four and two days, respectively. Treated cells were collected and RNA was extracted using the Qiagen (Hilden, Germany) RNA-mini kit or Norgen (Thorold, ON) single cell RNA purification kit.

#### Library preparation, sequencing, quality check, alignment, quantification of raw read counts

Library preparation, sequencing, quality check, alignment, quantification of raw read counts, and normalization of read counts were performed using the same methods as described in Mohammadnia et al. (2024) (23).

#### Analysis of RNA-seq data from human primary oligodendrocytes

Samples were analyzed in a pairwise manner as described in Mohammadnia et al. (2024) (23).

#### Analysis of RNA-seq data from iPSC-differentiated cells

As these samples did not exhibit significant heterogeneity and the lack of pair information for samples, we used DESeq2 for differential gene expression analysis following the methodology outlined in Pernin et al. (2024) (24, 25). Significantly differentially expressed genes were identified using a threshold of log2 fold change > 1 and a p-value cutoff of < 0.05.

Heatmaps were generated following hierarchical clustering using the “Hierarchical Clustering Image” function in GenePattern (v3.9.11). Gene clustering was performed using Pearson correlation, and normalized values were transformed using a log transformation. Finally, row normalization was applied for the final visualization (26). Single-sample Gene Set Enrichment Analysis (ssGSEA) was performed using the same method described in Mohammadnia et al. (2024)(23). However, the signatures used in this analysis were BMP4 target genes extracted from the weighted matrix of ligand-target interactions in NichNet (27). We selected target genes based on their weighted values, specifically those with a weight > 0.1 (n = 107 genes).

#### Single-cell RNA-seq analysis

All single-cell RNA-seq analyses were performed using the same pipeline employed in our previous publication (25) on the same Jakel et al. (2019) data set (28).

### Statistical analysis

Statistical analyses were performed using Excel or GraphPad Prism software. Cell culture studies were performed with at least three individual replicates per experiment. For iPSC-studies, three cell lines were used which contributed to values on graphs, and subsequent passages were used as biological replicates. Bars on graphs indicate means ±SEM. Student’s t-test was used for comparisons between two groups. P-values <0.05 were considered statistically significant.

## 3. Results

### Fibrinogen inhibits the differentiation of human OPCs

To corroborate previous findings in the field (11), we aimed to test the effects of fibrinogen on human oligodendrocyte progenitors. Since, as stated, we cannot isolate early OPCs from our surgical samples, we generated early OL-lineage cells from iPSCs. We also utilized a previously described human iPSC reporter line (SOX10mO) (20) where fluorescence intensity of the reporter correlates to the overall maturity of the cells. Fibrinogen treatment of early lineage cells had no significant effect on OPC viability (Fig. 1A), although treatment did significantly reduce the proportion of O4+ OPCs (Fig. 1B) as measured by immunofluorescence. To assess the effect of fibrinogen on overall differentiation into the OL lineage, we assessed level of our SOX10mO reporter. Under basal conditions, as shown in Fig. 1C, placing these cells in differentiation media (see Materials and Methods) results in increased SOX10mO expression with a reduced proportion of undifferentiated (SOX10-neg) cells. Addition of fibrinogen resulted in an increase proportion of SOX10-med cells. We used the astrocyte-specific marker CD49f (29) to follow-up on the observation by Petersen et al. (11) that fibrinogen diverts the differentiation of primary post-natal rodent derived OPCs towards the astrocyte lineage. We observed that the overall decrease in O4+ cells was associated with a significant increase in proportion of O4-CD49f+ cells and O4+CD49f+ cells following fibrinogen exposure (Fig. 1D, E). The latter hybrid OPCs retained the expression of our reporter (Fig. S1). To explore the molecular response of our SOX10mO cells, they were FACS-selected as previously described (20). Bulk RNAseq analysis indicates an overall decrease in signalling and differentiation occurring in these early OL-lineage cells, alongside significant upregulation of cell-cycle pathways (Fig. 1G-H), confirmed by Ki67 staining (Fig. S2). These findings coincide with findings by Petersen et al. (11) in primary rodent OPCs and serve as confirmation for fibrinogen acting on OPC differentiation capacity in a human context.

### Human primary OLs have an age and maturation-dependent functional response to fibrinogen

To further explore the effect of fibrinogen across the OL lineage, dissociated human primary OL cultures comprised of 95 % mature OLs (19), were used. Following treatment, we observed no significant effect on cell viability (Fig. 2A), regardless of added concentration of fibrinogen. To determine a functional effect on OLs, we assessed their capacity to ensheath poly-L-lactide nanofibers. We separated pre-myelinating OLs (A2B5+) from mature OLs (A2B5-) to determine the relation of lineage maturity. In the presence of fibrinogen, we observed an increased capacity in adult-derived mature OLs (A2B5-) (p < 0.05) but no effect on adult A2B5+ cells (Fig. 2B). Previous studies in our lab suggest that pediatric-derived primary A2B5+ cells most closely resemble a precursor (19). Therefore, to extend our studies across the entirety of the OL lineage, we next sought to determine the effect of fibrinogen on pediatric donor cells. Following exposure, we observed a significant decrease in ensheathing A2B5+ cells (p < 0.05) but not mature OLs (A2B5-) in comparison to the age-matched untreated controls (Fig. 2C). Figures 1D-E show the overall age-related effect of fibrinogen on mature (A2B5-) cells (Fig. 2E) and immature (A2B5+) cells (Fig. 2D), with representative images in Fig. 2F, indicating that fibrinogen has its most apparent positive effect on cells derived from older adult donors. To confirm our in vitro findings, we performed Bulk RNAseq analysis of adult OLs. The transcriptome of adult-derived mature OLs treated with fibrinogen shows upregulation of lipid metabolism pathways, including cholesterol synthesis, (Fig. 2G-H) consistent with this beneficial effect of fibrinogen on ensheathment by adult OLs.

**Figure 2.**
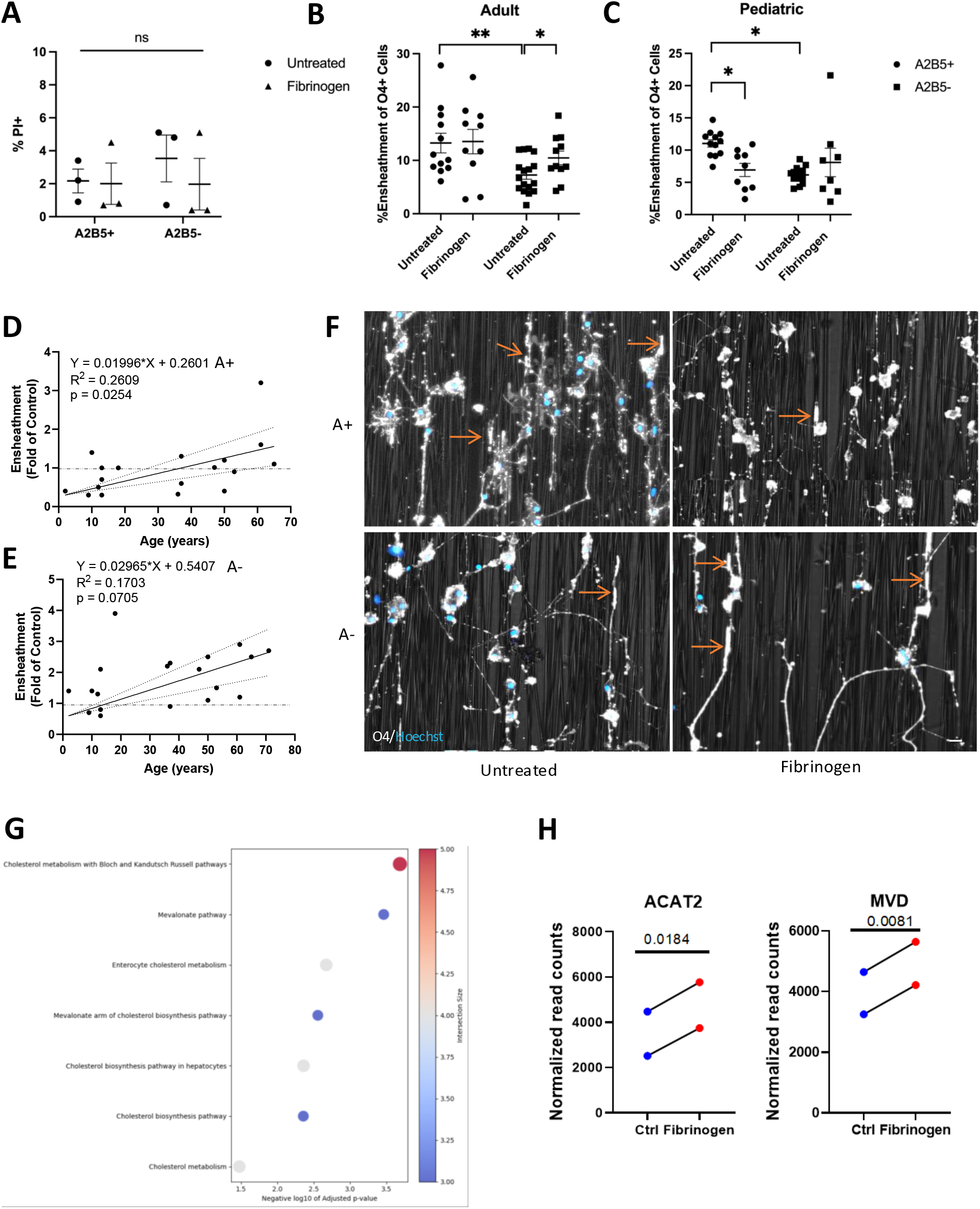
Fibrinogen has an age and maturation-dependent effect on human OLs. **A**. Cell viability of human A2B5+ pre-OLs (A+) and A2B5-mature OLs (A-) from both adult and pediatric samples following fibrinogen treatment and quantified following immunofluorescence microscopy for propidium iodide (PI). Bars indicate SEM. **B-C**. Capacity of (B) adult (26 - 65 years) and (C) pediatric (2.5 -18 years) pre-OLs (A2B5+ (A+)) and mature OLs (A2B5- (A-)) to ensheath nanofibers. * p < 0.05, ** p<0.01, *** p<0.001. Bars indicate SEM. Paired t-tests were applied to assess significance levels in all plots. **D-E**. Correlation of (D) A2B5+ and (E) A2B5-cell ensheathment capacity to age, as normalized to age-matched controls following treatment with fibrinogen. **F**. Representative images of adult OLs on nanofibers. O4 stain in white, nuclei in blue. Top panels represent A2B5+ cells, bottom panels represent A2B5-cells. Arrows point to representative examples of ensheathing segments. Representative images were cropped and brightness was increased to best show areas of interest. **G**. GO term analysis on upregulated genes following bulk RNA sequencing on fibrinogen-treated adult OL samples. **H**. Significantly upregulated genes involved in lipid metabolism, following fibrinogen treatment on adult OL samples (matched donors).

### Fibrinogen signalling in human OL lineage cells via the BMP pathway

Fibrinogen has been previously suggested to act via the BMP pathway in rodent OL lineage cells, and inhibiting this pathway abrogates its effect (11). To determine if the BMP pathway was activated in human cells throughout the OL lineage in response to fibrinogen, we measured the activation of a downstream signalling factor, SMAD1/5/9 (30). Nuclear localization of phosphorylated SMAD (pSMAD), as opposed to cytoplasmic, indicates activation, and we used this as a readout for our study. In mature OLs, we observed a significant increase in nuclear localization of pSMAD1/5/9 in response to fibrinogen (Fig. 3A-B) in comparison to untreated cells. To confirm pathway activation earlier in the OL lineage, human iPSC derived OPCs were exposed to fibrinogen in the same manner, and we a similar trend (Fig. 3C). Comparing the early OPCs and the mature OLs (Fig. 3E), nuclear pSMAD localization was more frequent in the latter, indicating that the BMP pathway is more activated in these cells than iPSC-derived cells after exposure to fibrinogen. For both primary adult and iPSC-derived O4+ cells, BMP4 treatment induced similar changes and fibrinogen-induced responses were significantly reduced by the BMP-pathway inhibitor, DHM1 (Fig. 3).

**Figure 3.**
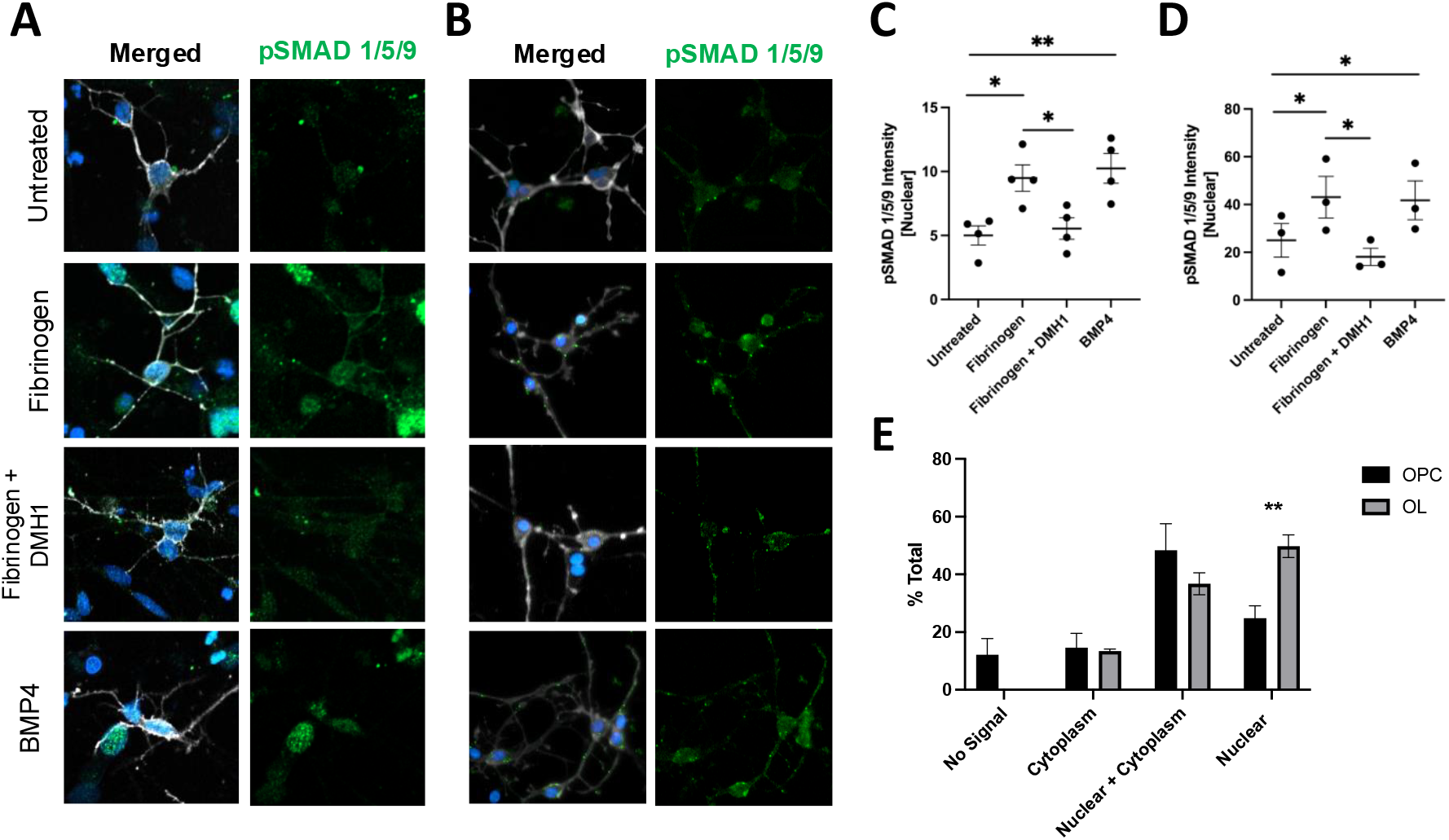
Fibrinogen activates BMP signalling across the OL lineage. **A-B**. Representative images of (A) human iPSC derived OPCs (O4+) and (B) human primary OLs (O4+) following exposure to treatments for 2 hours, by Zeiss confocal microscope. pSMAD 1/5/9 (green), O4 (white), nuclei (blue). Representative images were cropped and brightness was increased to best show areas of interest. **C-D**. Quantification of nuclear fluorescence intensity of pSMAD 1/5/9 in (C) iPSC OPCs and (D) human primary OLs, where fibrinogen and BMP4 significantly increase pSMAD1/5/9 nuclear signal intensity. **E**. Proportion of pSMAD signal localization following treatment with fibrinogen. n=3 donors for primary cells, n=4 passages for iPSC * p < 0.05, ** p<0.01, *** p<0.001. Bars indicate SEM. Paired t-tests were applied to assess significance levels in all plots.

Confirming these findings transcriptomically, Bulk RNA seq analyses of fibrinogen-treated mature OLs show activation of the BMP pathway (Fig. 4A), as well as an array of BMP-regulated genes (Fig 4C-D). Analysis was based on paired samples given the diversity between donors. In regards to the early OL-lineage cells, we detected a non-significant but decreasing trend in BMP pathway enrichment (Fig 4B); however, we did not observe significant changes in BMP regulated genes including those of the astrocyte lineage (data not shown). The early OL-lineage cells showed an overall gene suppression except for increase in cell proliferation related pathways (Fig. 2H-I) which could explain the lack of BMP-regulated genes observed. These data show, that although fibrinogen activates the BMP-pathway in vitro both in early and mature OL-lineage cells, the downstream genes which are activated remain different.

**Figure 4.**
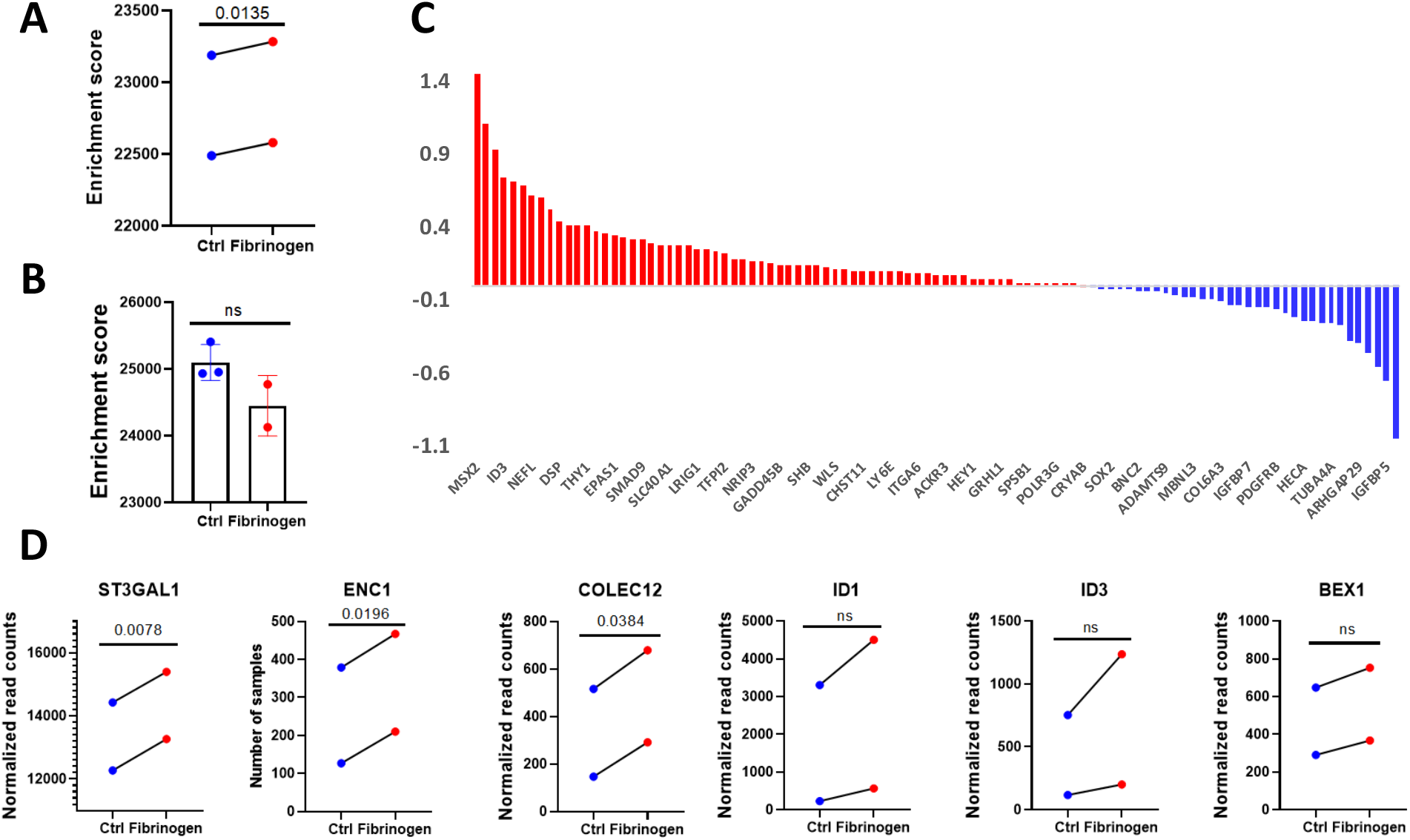
Transcriptomic analysis of BMP signaling activation in the oligodendrocyte lineage. **A-B**. Single-sample Gene Set Enrichment Analysis (ssGSEA) for BMP4 target genes in: (A) human primary OLs treated with fibrinogen at 2.5 mg/mL, 2 days and (B) iPSC-differentiated SOX10-positive medium cells treated with fibrinogen at 2.5 mg/mL for 4 days. Normalized read counts were used for both analyses. Enrichment scores are visualized in the plots. One iPSC-OPC fibrinogen-treated sample was removed during quality control. **C**. Bar plot showing log2-transformed fold changes (ratios) of BMP4 target gene expression in human primary OLs. Red bars indicate genes with higher expression levels in fibrinogen-treated cells, and blue bars indicate genes with higher expression levels in the control group. **D**. Example of BMP4 target genes in human primary OLs showing upregulation upon treatment with fibrinogen (2.5 mg/mL, 2 days). Paired t-tests were applied to assess significance levels in all plots for human primary OLs. In all plots, blue represents control samples, and red represents fibrinogen-treated samples.

### OLs in the MS brain parallel the response seen *in vitro*

To determine in-vivo relevance of our findings, we re-analyzed a publicly available RNAseq dataset from the human post mortem MS brain (28) (Fig 5A-C). Following re-clustering of the OLs, six sub-clusters were labelled Olig1-6, containing cells from both MS and non-neurologically diseased control samples. To confirm in vitro findings and explore BMP-pathway related signatures, each OL cluster was evaluated on the expression of top 107 target genes for the BMP4 ligand (Fig 5D). OL clusters 2-6 had more BMP genes expressed in cells derived from MS tissue rather than control. Distribution of known BMP-target genes *ID1* and *ID3* were evaluated and found to have an increasing trend in the cells derived from MS tissue (Fig 5E). To parallel this response to the signature found in our fibrinogen-treated adult OLs, the expression of lipid-related genes *ACAT2* and *MVD* was evaluated and found to have a similar increased trend in MS-derived samples (Fig 5F). These transcriptomic findings suggest that the beneficial effect of fibrinogen seen in vivo on mature OLs, may also be possible in the MS brain. In addition, our preliminary immunofluorescence studies of MS tissue show co-localization of fibrinogen with oligodendrocytes (see Supplementary Figure S3).

**Figure 5.**
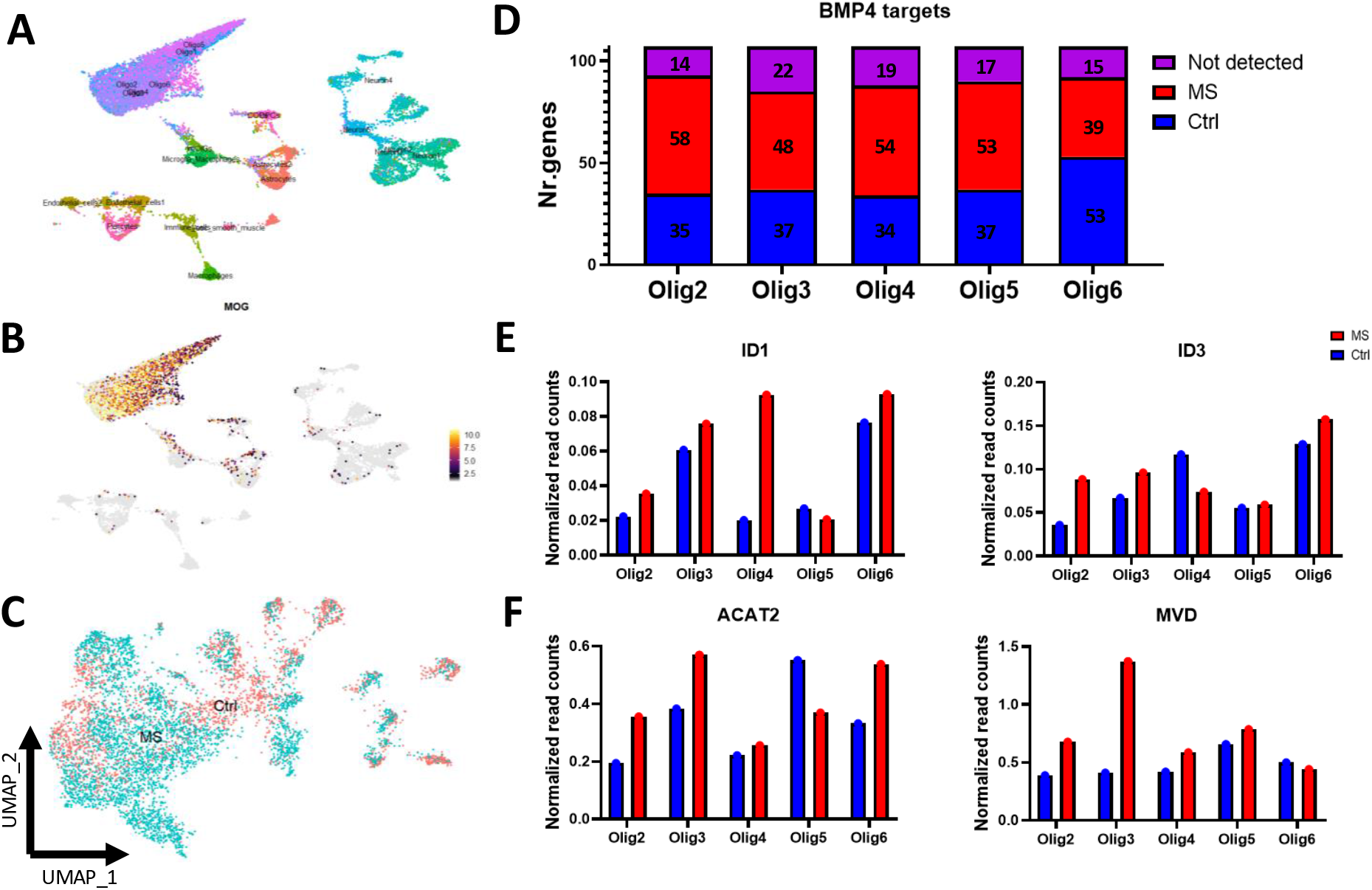
Transcriptomic analysis of the BMP signaling pathway in mature oligodendrocytes using single-nucleus RNA-sequencing (snRNA-seq) data. **A**. UMAP plots were generated based on the study by Jakel et al. (2019) to capture all relevant cell types in the context of multiple sclerosis (MS) and control tissue. **B**. Mature OL marker genes were used to identify mature cells within the snRNA-seq data. Myelin OL glycoprotein (MOG) is visualized in this UMAP plot as an example of an OL-specific marker gene. **C**. All cell clusters contained a mixed combination of cells from MS and control patients, making them suitable for comparing BMP signaling within each OL population. **D**. Bar plot showing the distribution of BMP4 target genes across different OL populations in MS and control groups. We used the top 107 target genes for the BMP4 ligand (with interaction weight > 0.1) and extracted their average expression in each OL cluster from both MS and control groups. The Oligo1 population was excluded because most of its cells (85%) originated from control samples. In the bar plot, purple, red, and blue represent not expressed, higher expression in MS, and higher expression in control samples, respectively. **E**. ID1 and ID3 are examples of BMP signaling target genes that show an increasing trend in expression in most OL populations in MS compared to control groups. **F**. Example of genes related to lipid metabolism that exhibit a similar increasing trend in gene expression as BMP signaling target genes, consistent with the results from *in vitro* experiments using human primary OLs. In all cases, normalized read count data were used to calculate average expression levels for each cluster for analysis and plot generation. Red represents MS, and blue represents control cells.

## 4. Discussion

Fibrinogen, a blood coagulation protein, is deposited in many CNS diseases with BBB disruption and myelin abnormalities, including MS, stroke and traumatic brain injury (13). Its involvement throughout the MS disease course is highlighted by studies showing disruption of the BBB and fibrin deposition early in MS lesions and preceding demyelination (31), while in progressive MS with severe cortical fibrinogen deposition there is reduced neuronal density (7).

Fibrinogen has been noted to have a functional effect on immune cells, and resident CNS glia such as astrocytes and microglia (13, 32, 33). This molecule has been tested on rodent OPCs and OLs (11), but, to our knowledge, no studies have documented the effect of fibrinogen on OL-lineage cells in human samples. Here we demonstrate the *in vitro* effect of fibrinogen on human OL-lineage cells is dependent on differentiation stage and preliminary evidence of co-localization of fibrinogen on OL-lineage cells in MS brain tissue (Fig S3).

As our primary cell isolation technique is not conducive to isolation of early lineage cells (PDGFR⍰+ A2B5+ O4+), we derived OPCs from human iPSCs and employed a previously characterized iPSC reporter line (SOX10mO) to aid in determining the overall culture maturation stage (20). We found reduced O4+ cells when fibrinogen was added during the last four days of differentiation, in line with previous rodent studies (11). This data is further compatible with the inhibition of ensheathment by pediatric A2B5+ cells, the most progenitor-like cells within our primary cell isolates. Using the SOX10mO line, we observed an increase in the proportion of SOX10-med cells. We relate this finding to increased proliferation in response to fibrinogen, as shown by Ki67 and bulk RNAseq, which matches the data observed in rodent OPCs (11). The decrease in proportion of O4+ cells was associated with the increase in cells co-expressing the astrocyte surface marker, CD49f. This deviation towards the astrocyte lineage parallels with the findings of Petersen et al. (11), although was not evident in our bulk RNAseq data. This may be due to this being a small sub-population of cells less prominent within the larger SOX10-medium sample, where overall cell cycling genes were more predominantly upregulated.

The mature OL plays important roles in myelin maintenance and can even contribute to re-myelination in the post-natal brain (34). To assess the functional effects of fibrinogen on OLs in relation to their stage in the cell lineage, we obtained mature human OLs from surgically resected materials as described previously (17, 18). These samples were characterized further into late progenitors (A2B5+) and mature OLs (A2B5-) both from pediatric and adult donors. Our functional studies indicate a differential effect of fibrinogen in relation to OL lineage stage. Mature OLs (A2B5-), most apparent with adult donor cells, had increased ensheathment capacity following fibrinogen exposure, whereas fibrinogen had detrimental effects on ensheathment capacity of pediatric A2B5+ cells. Bulk RNAseq supported these positive observed effects in adult cells, showing enrichment for lipid metabolism pathways as would be upregulated in OL-mediated ensheathment and subsequent remyelination (35, 36). Pediatric and adult human OL-lineage cells differ on a transcriptomic level, with the indication that pediatric cells are more progenitor-like (19). These results indicate that both donor age and lineage stage underlie functional differences.

The work of the Akassoglou group determined that the effects of fibrinogen on OL-lineage cells are mediated by signaling via the BMP pathway. Using postnatal rat O4-selected cells (11), they confirmed this with a knockout model of the BMP type-1 receptor molecule, activin A receptor type I (ACVR1). Our analysis of this pathway indicates differences between the primary and the iPSC-derived OL-lineage cells. Using immunocytochemistry, we observed a strong nuclear translocation of the downstream BMP effector molecule, pSMAD 1/5/9 in mature OLs; this response was less marked in the iPSC-derived cells. For both groups, we showed fibrinogen signaling inhibition using the ACVR1 inhibitor DMH1, showing there is some functional signaling. Our RNAseq data provides some insight into the basis of the observed functional responses. Extensive gene upregulation downstream of the BMP pathway was observed in the adult primary OLs, which was paralleled in a subset of OLs from an MS RNAseq dataset and coincided with upregulation of lipid synthesis pathways in both data sources. On the other hand, the RNAseq on early OL-lineage cells showed a downregulation of all signaling pathways in response to fibrinogen, and as a result there was no enrichment for BMP pathway associated genes.

As our results have shown that the effects of BMP pathway activation are highly context dependent. In the perspective of development, studies have determined that BMP signaling has different effects dependent on developmental stage such as promoting neural progenitor cells to proliferate in early development while promoting astrocytic and neuronal differentiation in later stages (37). These previous findings in the literature may suggest a reason for our observed age-related effect of fibrinogen in human primary OLs. As seen in our iPSC-OPCs, BMP4 signaling also has the capacity to retain cells in a progenitor-like state (38), and promotes a subset of cells to gain an astrocytic phenotype. These observed differences can be attributed to ligand and receptor availability, as well as the presence of competing signaling molecules (39), although this would require further exploration in our model.

In the context of MS, the fibrinogen-driven increased ensheathment by mature OLs could positively contribute to the remyelination process that occurs during early inflammatory demyelinating activity with ongoing breakdown of the BBB, and at the borders of chronic active plaques. In these latter situations numbers of mature OLs are relatively preserved. Failure of subsequent remyelination would reflect a depletion of such cells, as evident in previously demyelinated regions of the plaque, that is not replenished by production of new OLs capable of remyelination due a block in oligodendroglial progenitor differentiation. This could reflect at least in part the effect of fibrinogen on OPCs. Whether other factors in the milieu of the MS lesion contribute to this differentiation block remains to be determined.

## Conflict of Interest

The authors have nothing to disclose.

## Author Contributions

G.J.B., Q.L.C., C.W., J.P.A and G.R.W.M. contributed substantially to the conception and design of the study. G.J.B., J.P.A., G.R.W.M. and A.M. drafted a significant portion of the manuscript or figures. G.J.B., Q.L.C., C.W., A.G., A.M.R.G., C.P., A.T., L.C.M., M.Y., E.C., J.S., V.E.C.P., J.A.H., R.W.R.D., M.S., S.E.J.Z., W.K., S.L., A.P., T.M.D., J.A.S. contributed to acquisition and analysis of data.

## Acknowledgements

We thank lab members Lama Fawaz, Vanessa Omana, Wolfgang Reintsch, Andrea Krahn, and Genevieve Dorval for technical or administrative assistance. The flow cytometry work and cell sorting were performed at the MNI’s Early Drug Discovery Unit’s Flow Cytometry Core Facility. We acknowledge the McGill microscopy platform for their support. This project was supported in part by a grant from MS Canada.

## Data Availability

data will be made available upon manuscript acceptance.

## Notes

### Competing Interest Statement

The authors have declared no competing interest.

### Summary of Updates

Figures have been rearranged to enhance clarity of the manuscript. Figure 6 has been put as a supplemental file.

