## Supplemental Files for "Age and maturation stage linked consequences of fibrinogen on human oligodendroglia"

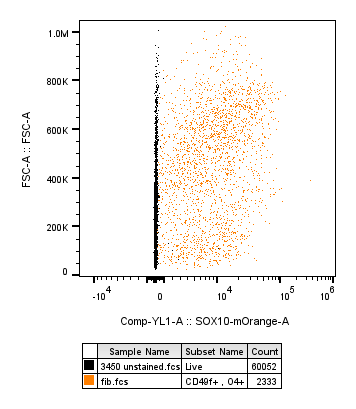

**Figure S1. mOrange signal in hybrid OPCs.** Following 4-day treatment of fibrinogen, data acquired by flow cytometry. CD49f+O4+ cells express mOrange (orange) at a higher fluorescence intensity in comparison to non-reporter cells (black).

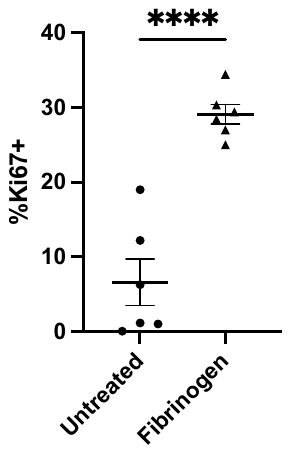

**Figure S2. Proliferation in fibrinogen treated OPC cultures.** Proportion of Ki67+ nuclei quantified in iPSC-OPC cultures treated with fibrinogen. p < 0.0001. n=6 replicates, bars indicate SEM.

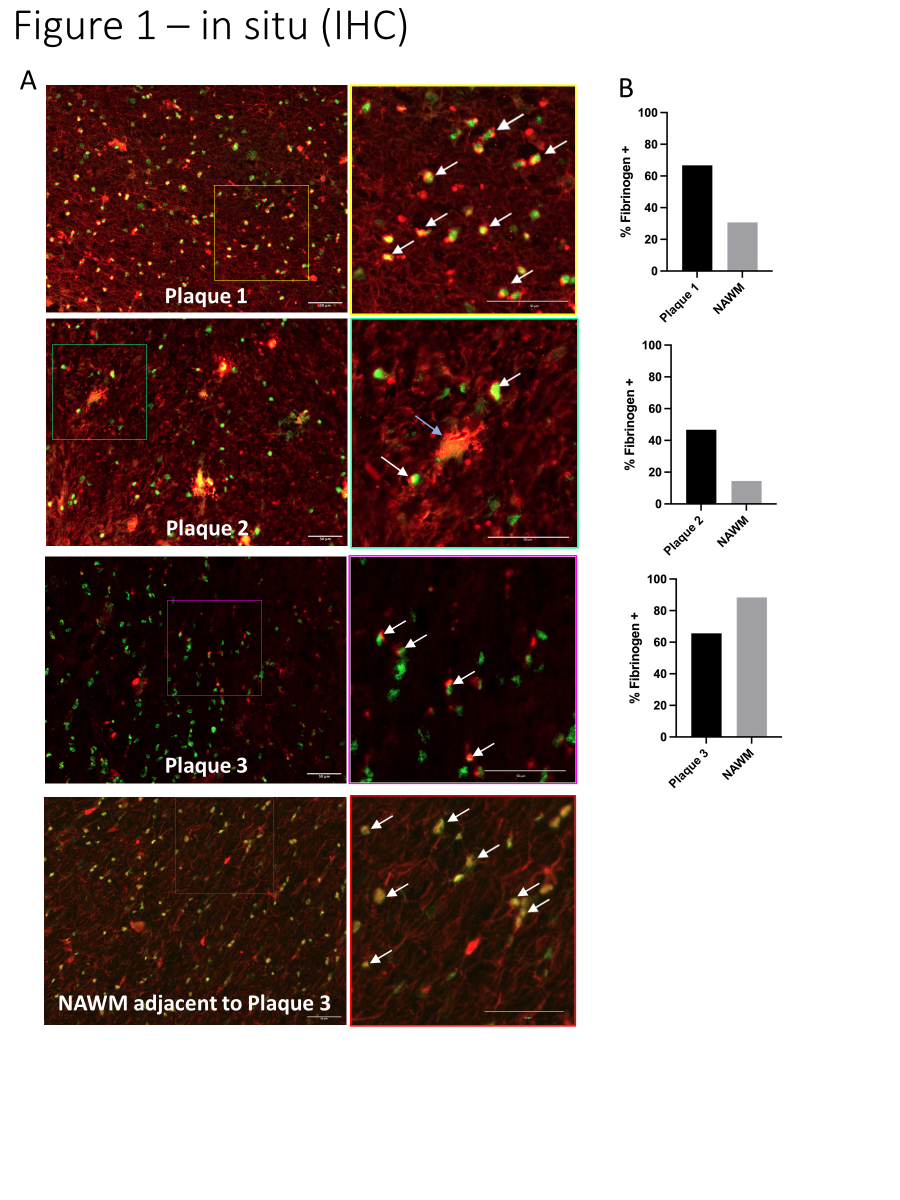
**Figure S3. Colocalization of fibrinogen and OL lineage cells in the MS brain.**

Representative images of MS plaque and MS NAWM. Sox10 (oligodendroglial nuclear marker) in green; fibrinogen in red. For each region, a higher magnification of the area within the box is shown in the panel immediately to the right. Plaque 1, an MS lesion that in sections stained for myelin showed active demyelination at its edge, consistent with a chronic active plaque; Plaque 2, an MS lesion that in sections stained for myelin showed no active ongoing demyelination, consistent with a chronic inactive plaque. Plaque 3, an MS lesion that in sections stained for myelin showed active demyelination throughout its extent, consistent with an active plaque. The representative NAWM illustrated is adjacent to Plaque 3. Examples of SOX10-positive oligodendroglia that are positive for fibrinogen in their cytoplasm and/or nuclei are indicated with white arrows. The blue arrow indicates a prominent fibrinogen-positive cell, possibly an astrocyte or macrophage that has engulfed SOX10- and fibrinogen-positive debris: the debris double labelled with red and green, appearing yellow. Scale bars = 50µm.

**Methodology**

**Human MS brain tissue**

Tissues from the Centre Hospitalier de l’Université de Montréal (CHUM) were obtained by a Material Transfer Agreement between CHUM and McGill University. Human brain tissue was obtained from patients diagnosed with MS according to the revised 2010 McDonald’s criteria (1). Tissue samples were collected from three MS patients (Table S1) with full ethical approval (BH07.001, Nagano 20.332-YP) and informed consent as approved by the local ethics committee. Autopsy samples were preserved, and lesions classified (2) using Luxol Fast Blue-Haematoxylin and Eosin (LFB-HE) staining as previously published (3, 4).

**Immunohistochemistry**

Frozen human brain tissue sections, previously briefly fixed in acetone, were further fixed in 100% acetone for 10 minutes, followed by 70% ethanol for 5 minutes. Subsequently, sections were washed with PBS and 0.05% PBST (phosphate-buffered saline + Tween20) for 3 minutes. Fc-receptors in sections were blocked with PBS containing 10% donkey serum. Primary antibodies, goat anti-SOX10 (R&D systems, Oakville ON, AF2864) and rabbit anti-fibrinogen (Agilent, Santa Clara CA, A0080) were diluted 1:500 (0.2 mg/ml) and 1:400 (1mg/ml) respectively in PBS containing 3% donkey serum and incubated overnight at 4℃. Isotype controls consisted of the IgG of the same species and same concentration as each primary antibody. Following incubation, sections were washed with 0.05% PBST 3 times for 5 minutes each. Fluorophore-conjugated secondary antibodies (Invitrogen, Waltham MA) were prepared at a 1:200 dilution in PBS, donkey anti-goat IgG Alexa Fluor 488 to visualize SOX10 and donkey anti-rabbit IgG Alexa Fluor 647 to visualize fibrinogen and slides were incubated in the dark for 60 minutes at room temperature. Sections were washed with 0.05% PBST 3 times for 3 minutes and incubated with Hoechst (Invitrogen, Waltham MA; 1:5000) for 10 minutes at room temperature. Once washed with 0.05% PBST 3 times for 3 minutes to block lipofuscin autofluorescence, sections were incubated with 0.3% TrueBlack (Biotium, Fremont, CA) diluted in 70% ethanol for 3 minutes. Slides were washed with PBS and mounted with PermaFluor mounting medium (Thermo-Fisher, Missisauga, ON), coverslipped and stored in the dark at 4℃.

1. Polman CH, Reingold SC, Banwell B, Clanet M, Cohen JA, Filippi M, et al. Diagnostic criteria for multiple sclerosis: 2010 revisions to the McDonald criteria. Ann Neurol. 2011;69(2):292-302.
2. Kuhlmann T, Ludwin S, Prat A, Antel J, Brück W, Lassmann H. An updated histological classification system for multiple sclerosis lesions. Acta Neuropathol. 2017;133(1):13-24.
3. Dhaeze T, Tremblay L, Lachance C, Peelen E, Zandee S, Grasmuck C, et al. CD70 defines a subset of proinflammatory and CNS-pathogenic T(H)1/T(H)17 lymphocytes and is overexpressed in multiple sclerosis. Cell Mol Immunol. 2019;16(7):652-65.
4. Broux B, Zandee S, Gowing E, Charabati M, Lécuyer MA, Tastet O, et al. Interleukin-26, preferentially produced by T(H)17 lymphocytes, regulates CNS barrier function. Neurol Neuroimmunol Neuroinflamm. 2020;7(6).

**Table S1. Tissue Donor information.**

| **Section** | **Age**  **(years)** | **Sex** | **Disease Type** | **EDSS** | **Disease Duration** | **Post-mortem**  **interval** |
| --- | --- | --- | --- | --- | --- | --- |
| MS Plaque 1 | 48 | M | SPMS | 8.5 | 6 years | N/A |
| MS Plaque 2 | 60 | F | SPMS | 8.5 | 28 years | 2.5 hrs |
| MS Plaque 3 | 44 | F | RRMS | 7 | 13 years | 5 hr |

Abbreviations: EDSS=Expanded Disability Status Scale, F= Female, M=Male, MS=Multiple Sclerosis, N/A=Not Available, RRMS=Relapsing-remitting MS, SPMS=Secondary Progressive MS.

**Table S2. iPSC Line Information**

| **Cell Line Name** | **Sex** | **Patient Age (years)** | **Material Source** | **Type of Reprogramming** |
| --- | --- | --- | --- | --- |
| AIW002-02 | M | 37 | PBMC | Retrovirus |
| 3450 | M | 37 | PBMC | Episomal |
| SOX10mOrange (3450) | M | 37 | PBMC | Episomal |

Abbreviations: M=Male, PBMC= Peripheral Blood Mononuclear Cells.

**Table S3. Antibodies and Reagents**

| **Antibody** | **Supplier** | **Identifier** |
| --- | --- | --- |
| goat anti-SOX10 | R&D systems, Oakville ON | AF2864 |
| rabbit anti-Fibrinogen | Agilent, Santa Clara CA | A0080 |
| isotype control polyclonal rabbit | BioLegend, San Diego CA | 910801 |
| isotype control goat IgG | R&D systems, Oakville ON | AB 108-C |
| O4-APC | Miltenyi, Auburn CA | 130-119-982 |
| Human TruStain FcX Fc Receptor Blocking Solution | BioLegend, San Diego CA | 422302 |
| LIVE/DEAD Fixable Aqua Dead Cell Stain Kit, for 405nm excitation | Invitrogen, Waltham MA | L34957 |
| CD49f-PE-Dazzle | BioLegend, San Diego CA | 313626 |
| mouse IgM anti-O4 | R&D, Oakville ON | MAB1326 |
| rabbit anti-pSMAD1/5/9 | Cell Signalling Technology, Danvers MA | 13820S |
| rat anti-Ki67 | Invitrogen, Waltham MA | 14-5698-82 |
